# Mutational analysis and assessment of its impact on proteins of SARS-CoV-2 genomes from India

**DOI:** 10.1101/2020.10.19.345066

**Authors:** Rezwanuzzaman Laskar, Safdar Ali

**Author notes:** **Corresponding author:** Dr Safdar Ali, Assistant Professor, Department of Biological Sciences, Aliah University, IIA/27, Newtown, Kolkata 700160, India.

## Abstract

The ongoing global pandemic of SARS-CoV-2 implies a corresponding accumulation of mutations. Herein the mutational status of 611 genomes from India along with their impact on proteins was ascertained. After excluding gaps and ambiguous sequences, a total of 493 variable sites (152 parsimony informative and 341 singleton) were observed. The most prevalent reference nucleotide was C (209) and substituted one was T (293). NSP3 had the highest incidence of 101 sites followed by S protein (74 sites), NSP12b (43 sites) and ORF3a (31 sites). The average number of mutations per sample for males and females was 2.56 and 2.88 respectively suggesting a higher contribution of mutations from females. Non-uniform geographical distribution of mutations implied by Odisha (30 samples, 109 mutations) and Tamil Nadu (31 samples, 40 mutations) suggests that sequences in some regions are mutating faster than others. There were 281 mutations (198 ‘Neutral’ and 83 ‘Disease’) affecting amino acid sequence. NSP13 has a maximum of 14 ‘Disease’ variants followed by S protein and ORF3a with 13 each. Further, constitution of ‘Disease’ mutations in genomes from asymptomatic people was mere 11% but those from deceased patients was over three folds higher at 38% indicating contribution of these mutations to the pathophysiology of the SARS-CoV-2.

## Introduction

The ongoing COVID-19 global pandemic began from Wuhan, China and has devastated millions of lives, economies and even nations as a whole. The first reported case was in December 2019 and as of 1^st^ September 2020, there have been 2,56,21,967 reported cases and 8,54,235 deaths worldwide (www.worldometers.info/coronavirus/). Of these, 36,21,245 cases and 64,469 deaths have happened in India making it one of the most affected countries in the world (www.mygov.in/covid-19).

The causative agent identified for COVID-19 is Severe Acute Respiratory Syndrome Coronavirus 2 (SARS-CoV-2) which belongs to family *Coronaviridae* characterized by single strand positive sense RNA genome. Though this is novel virus but the outbreak is not the first one from members of *Coronaviridae.* Previously, severe acute respiratory syndrome (SARS) coronavirus in 2002 and Middle East respiratory syndrome (MERS) coronavirus in 2012 created a scare as they had a relatively higher mortality rate. However, SARS-CoV-2 is by far the most contagious one [1–4].

The higher incidence of viral infections would imply a faster evolution process for SARS-CoV-2 [5]. This is so because more the virus replicates, higher are the chances of it accumulating mutations with the possibility of it leading to altered dynamics of its virulence, pathogenesis and interactions with host. The changes may not be necessarily favoring the virus; however, the unpredictability demands caution.

The SARS-CoV-2 genome encodes for 16 non-structural proteins in addition to the replicase polyprotein, the spike (S) glycoprotein, envelope (E), membrane (M), nucleocapsid (N) and other accessory proteins [6]. The impact of mutations in all the regions of the genome needs to be assessed to understand viral evolution.

With a definitive possibility of India becoming the most affected country by SARS-CoV-2 in near future and the demographic burden involved, its pertinent to be analyze the accumulating variations in the genome accounting for possible changes in protein and their potential to alter the virus in any manner. On 6th June, 2020 we retrieved 611 FASTA sequence congregations from India along their rational meta data from GISAID EpiCoV server to construct the phylo-geo-network and analyze the haplogroups along with their geographical distribution across different states of India [7]. Herein we extend our study using the same congregation of sequences to analyze the nature and composition of the observed mutations and their impact on proteins of SARS-CoV-2.

## Methodology

### Sequence Congregations Collection

GISAID EpiCoV is an open access repository of genomic and epidemiologic information about novel corona viruses from across the world from wherein sequences were extracted and alignment performed as previously reported [7]. Briefly, 611 FASTA sequence congregations along their rational meta data from GISAID EpiCoV (www.epicov.org) server. For mutational profile analysis with clinical correlation, we selected 15 genomes of deceased patients from existing congregation. However, there were just two genomes for asymptomatic patients in the congregations. So, on 09.12.2020, we downloaded 775 FASTA sequences with patient status from the same server and selected 30 genomes from asymptomatic patients. As the data filter for genome extraction, we used hCoV-19 as a virus name, human as a host, India as a location and complete sequence with high coverage. Details of the asymptomatic samples are given in Supplementary file 1. The sequences thus extracted were analyzed with NC_045512.2 from Wuhan, China as reference.

### Nucleotide Analysis

MEGA(v.10) is a multithreaded tool for molecular and evolutionary analysis. Multiple Sequence Alignment (MSA) of the extracted sequences [7] was initially visualized by this software then the variable sites are exported into spreadsheets with or without missing/ambiguous and gap sites along their respective positions [8]. Using this software, we estimate the MCL (Maximum Composite Likelihood) nucleotide substitution pattern and Tajima’s Neutrality test to understand transition transversion bias and nucleotide diversity [9, 10]. PIRO IGLSF is a MATLAB-based simulation software, we used this for the identify the location of mutated nucleotide position on specific gene [11].

### Protein Analysis

Coronavirus Typing Tool of Genome Detective (v.1.13) and COVID-19 Genome Annotator of Coronapp are webtools for analysis of protein and nucleotide mutation [12, 13]. We used these tools for annotation, identification and classification of mutated protein followed by verification and validation of the positions with the mutated nucleotide sites by the output of MEGA. The nucleotide similarity percentage was validated by NCBI BLAST (blast.ncbi.nlm.nih.gov) to investigate the sequence diversity.

### Protein Prediction Analysis

SIFT, PROVEAN and ws-SNPs&GO are the prediction tools which report positive or negative impact of variants on protein phenotype. The assessments are focused upon scores using several algorithms. It is expected that a SIFT score of < 0.05 is diseased (“affect protein function”), and that > 0.05 is neutral (“tolerated”). This is stated that a PROVEAN score of < −2.5 is diseased (“deleterious”), and > −2.5 is neutral. ws-SNPs&GO ’s PHD-SNP method is estimated to be > 0.5 mutation in the probability of disease, and < 0.5 is neutral [14–16].

## Results and Discussion

### Composition and distribution of variable sites

The observed MSA length was 29903 bp wherein the variable sites could be extracted through two different options. If we included gaps and ambiguous sequences, a total of 841 variable sites were observed with a percentage cover of 2.81%, where percentage cover = [(No of variable site / MSA Length) * 100]. All the sites have been shown in Supplementary file 2. The Tajima’s D statistic was also analysed (Table 1) and its negative value indicated the significance of these variable sites.

**Table 1:** Tajima’s Neutrality Test

| *m* | *S* | *Ps* | $\Theta$ | $\pi$ | *D* |
| --- | --- | --- | --- | --- | --- |
| 612 | 841 | 0.028124268 | 0.004021699 | 0.000412328 | -2.69529519 |

*m* = number of sequences, *n* = total number of sites, *S* = Number of segregating sites, *ps* = *S*/*n*, $\Theta$ = *ps*/*a*1, $\pi$ = nucleotide diversity, *D*= Tajima test statistic

However, excluding the gaps and ambiguous sequences reduced this percentage cover to 1.65% encompassing 493 variable sites which we have used for subsequent analyses reported in this study. This included 152 parsimony informative (PI) sites and 341 singleton sites (SNP: Single nucleotide Polymorphism). The PI sites are those whose incidence was observed in multiple samples whereas singleton sites had a restricted single sample incidence. The distribution of these sites according to various substitutions, protein localizations and impact therein has been summarized in Figure 1, Supplementary file 2. As evident therein, C→T (181 sites) forms the most prevalent mutation in both PI and singleton sites and G→T (95 sites) comes a distant second. The common aspect of two most prevalent mutations is “T” being the substituted nucleotide. Further, there were two multi-variable (MV) sites each in PI and singleton category wherein two separate mutations were observed at the same site in different samples. The details of observed MV sites have been summarized in Table 2.

**Figure 1:**
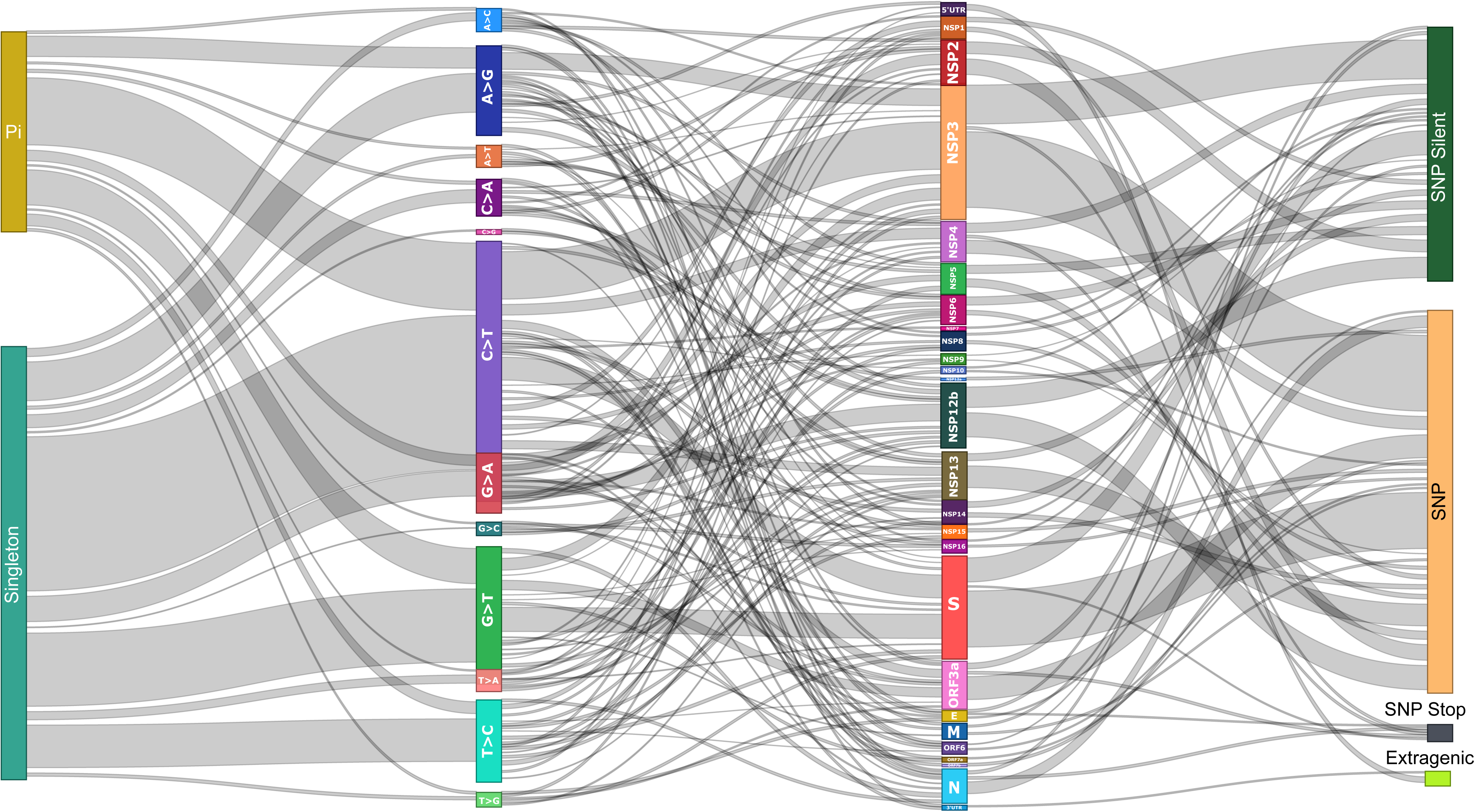
Summary of variations observed in SARS-CoV-2 genomes from India. The nature of variations (Singleton/PI); type of mutation, genome localization and impact on protein (SNP, SNP-Silent, SNP-Stop, Extragenic) has been represented along with their interlinking. The width of the connecting lines represent number; broader the line more the number of that parameter.

**Table 2:** Localization and mutations observed at the multi-variable (MV) sites

| S No | Nucleotide Position | Category | Nucleotide Variant 1 | Amino Acid Variant 1 | Nucleotide Variant 2 | Amino Acid Variant 2 |
| --- | --- | --- | --- | --- | --- | --- |
| 1 | 4893 | Pi | C>A | T725K | C>T | T725I |
| 2 | 5821 | SNP | A>G | L1034L | A>T | L1034F |
| 3 | 23282 | SNP | G>C | D574H | G>T | D574Y |
| 4 | 23593 | Pi | G>T | Q677H | G>C | Q677H |

The distribution of the variable sites across proteins of SARS-CoV-2 in a non-uniform manner is reflective of the differential contributions of proteins in evolution. As per our data, NSP3 had the maximum of 101 variable sites followed by S protein (74 sites), NSP12b (43 sites) and ORF3a (31 sites) (Figure 1; Supplementary file 2). These four proteins account for over half of the total variable sites of the genome and may be considered as drivers of genomic evolution for SARS-CoV-2. The mutations of S protein have been the focus for multiple research groups owing to its plausible impact on viral entry to the host cell but the mutations elsewhere may be equally relevant as the viral genome is known to harbor only what’s essential [17–19]. We believe a holistic approach is required to understand the evolution as more often than not the selection advantage being offered by any mutation is a chance event and can be from any part of the genome.

In terms of the impact of these variable sites on amino acid sequence of the viral proteins we classified them into four categories. First, the sites located in the extragenic region and hence no influence on the coding proteins. There were 10 such variable sites localized to the UTR regions (2 in 3’UTR and 8 in 5’UTR). Secondly, SNP-silent included those variable sites wherein the nucleotide change was leaving the amino acid sequenced unaltered. A total of 186 such sites were distributed across the genome. Thirdly, the variable sites which were leading to the introduction of a stop codon were referred to as SNP-stop and there were 8 such sites in our study. Lastly, the variable sites which were affecting the protein sequence are referred as SNP in the study and there were 281 such sites (Supplementary file 3). The prevalence and distribution of these sites has been summarized in Figure 1 and results of the prediction of their impact on protein has been discussed later.

### Constitution of genome, variable sites and the substitutions

In order to understand the underlying dynamics of substitutions, we performed the maximum composite likelihood estimate of nucleotide substitution as shown in Table 3. The reference and substituted nucleotide have been shown in rows and columns respectively. The values therein represent the probability of substitution (r) from one base to another. For simplicity, the sum of r values is made equal to 100. The nucleotide frequencies of the MSA are 29.87% (A), 32.14% (T/U), 18.37% (C), and 19.63% (G). The transition/transversion rate ratios are k1 = 2.195 (purines) and k2 = 7.799 (pyrimidines). The overall transition/transversion bias is R = 2.356, where R = [A*G*k1 + T*C*k2]/[(A+G)*(T+C)]. This substantiates the prevalence of certain mutations (C→T and G→T) over others.

**Table 3:** Maximum Composite Likelihood Estimate of Nucleotide Substitution. Rates of transitional substitutions are bold and transversional substitutions are *italicized*.

| Reference | Substituted |  |  |  |  |
| --- | --- | --- | --- | --- | --- |
|  | Nt | A | T | C | G |
|  | A | - | *4.57* | *2.61* | **6.13** |
|  | T | *4.25* | - | **20.39** | *2.79* |
|  | C | *4.25* | **35.68** | - | *2.79* |
|  | G | **9.33** | *4.57* | *2.61* | - |

We thereon looked at these variations in combination with their prevalence across samples. The most prevalent nucleotide at the variable sites in reference sequence was C (209) followed by G (137) whereas T was by far the predominantly substituted nucleotide (293, 60%). Also, the other three nucleotides had an almost equal representation in substitutions (A-68, G-68, T-64). This biased prevalence was not restricted to the alignment but was also getting translated to population incidence. There was a total of 723 mutations with C as reference nucleotide and 1032 mutations with T as substituted nucleotide across 611 studied genomes. The composition of 493 variable sites, their substitutions and prevalence across samples has been summarized in Figure 2 and Supplementary file 2. Evidently, any particular mutation may be incident across multiple samples and a single sample can harbor multiple mutations. A cumulative number for the same has been referred to as “Sum of Mutation Incidence” herein and thereafter in this study.

**Figure 2:**
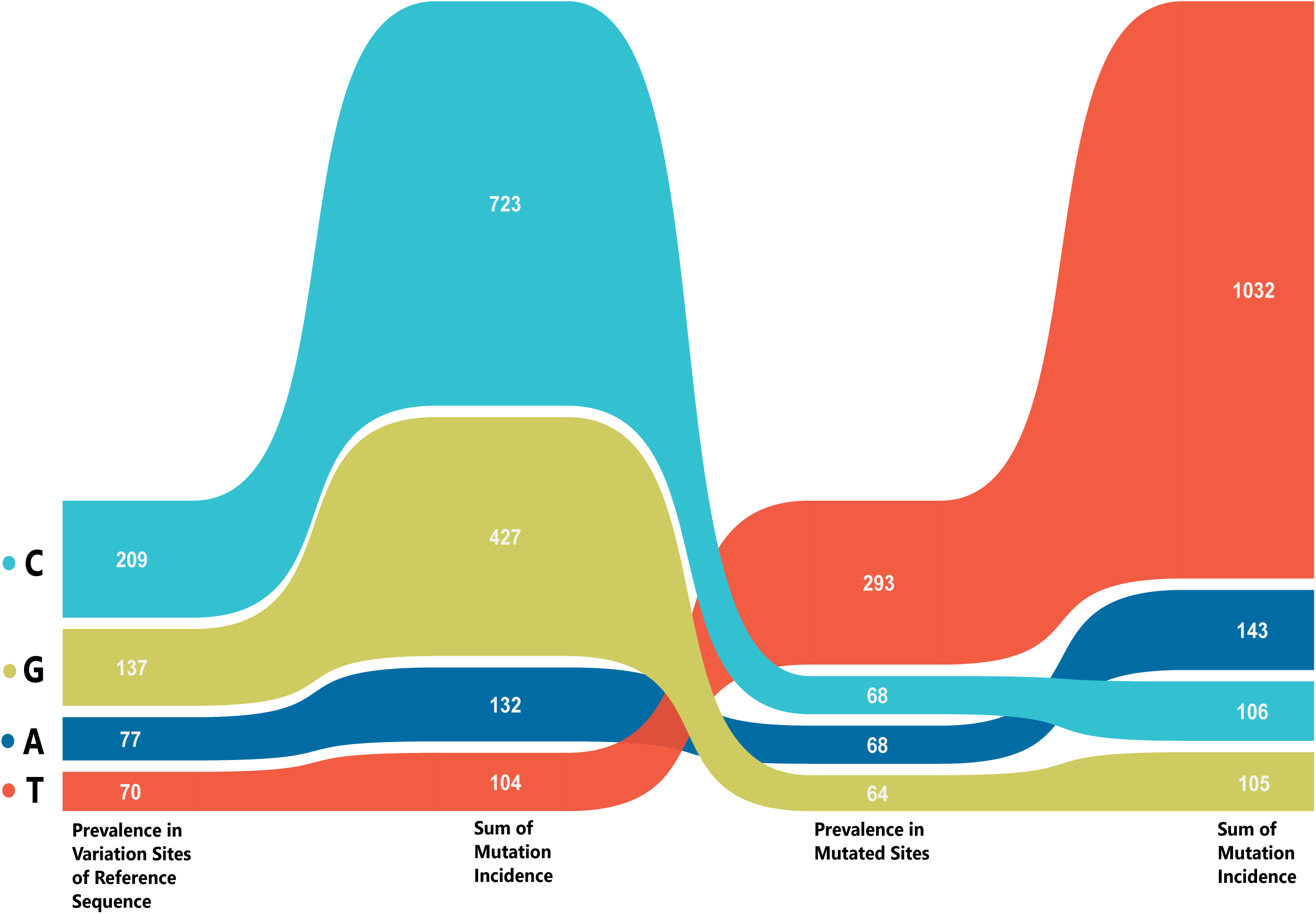
Prevalence and composition of different nucleotides across reference and substituted positions in SARS-CoV-2 genomes from India.

### Age and Gender wise distribution of samples and mutations therein

We subsequently analyzed the patient’s dataset with reference to age and gender for the incidence of mutations. However, since patients’ data wasn’t cumulatively available, the data for this aspect isn’t exhaustive but representative for 255 samples (104 females and 151 males). The patients whose genomes were used in the study and age was known were classified into seven categories from infancy to over 75 years. The maximum number of patients for both males and females belonged to mature adulthood category of 50 to 74 years with 57 and 40 samples respectively (Figure 3, Supplementary file 4). This adheres to the fact that the older population is at a greater risk for infection owing to a possibly weaker immune system and other physiological conditions.

**Figure 3:**
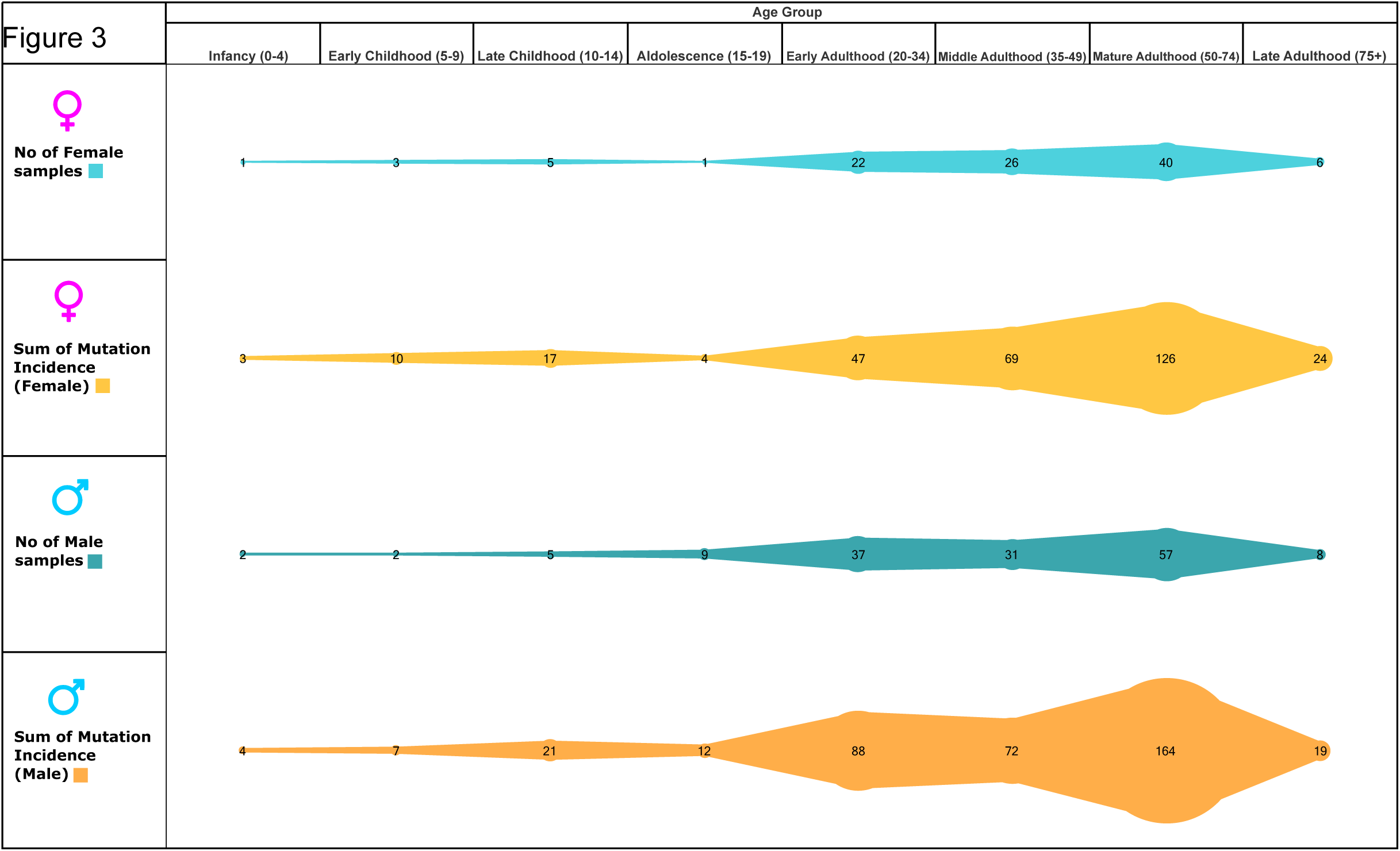
Age and gender wise distribution of mutations in SARS-CoV-2 genomes from India. Number of male/female samples and sum of mutations incidence therein according to age group.

The simple question of whether or not age and gender are associated with accumulation of genome variations has a not so simple answer. The overall average number of mutations per sample was 2.69 and the corresponding values for males and females separately was 2.56 and 2.88 respectively. Thus, women were contributing more to the mutational accumulation as compared to males. The individual mutational load for different age groups in males and females has been represented in Figure 4. Evidently, women are contributing more to the mutational load except for three age groups; 5-9years, 10-14years and 20-34 years. The highest difference on the basis of gender is for 15-19 years (2.67) but since there was just one female sample in that age group, it can’t be emphasized much in isolation but the overall pattern does seem relevant. This is more so because, in terms of incidence, males are almost 1.5 times of the females but in terms of variations, fewer females are contributing more to the mutational load. Possibly, the virus is behaving differently depending on gender.

**Figure 4:**
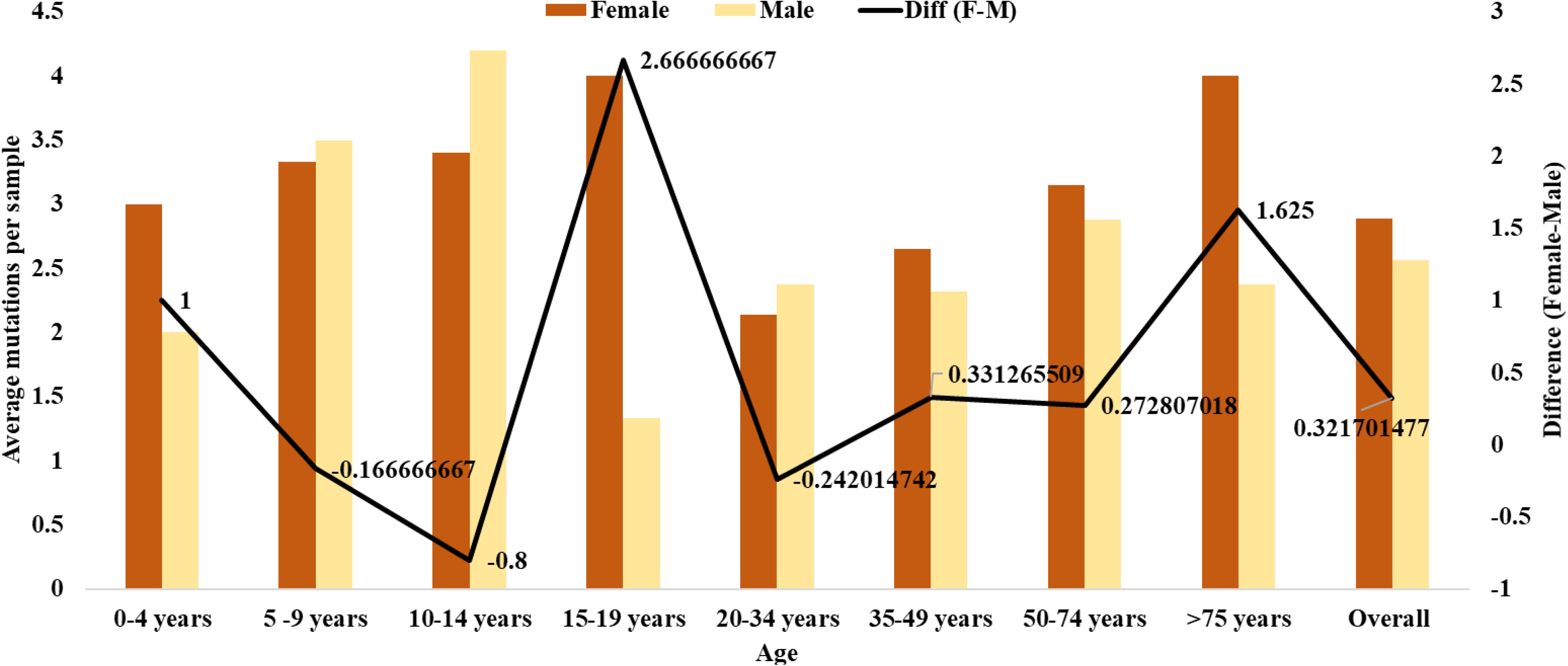
Average number of mutations per sample in different age groups of males/females and the differences therein.

### Geographical distribution and accumulation of variable sites

The mutational distribution across different states of India was subsequently ascertained. Generally speaking, more the virus replicates more should be the accumulated variations. The fact that the samples used in the study aren’t uniformly distributed across states provides for an intriguing template for analysis. The number of samples and the mutations therein for different states has been summarized in Figure 5 and Supplementary file 4. Evidently, Gujarat with highest number of 199 samples had maximum of 694 mutations. However, the correlation is neither uniform nor universal. For instance, Maharashtra with 94 samples had 203 variations whereas Telangana with 97 samples had 154 variations. A plausible explanation for this can be one sequence in Telangana (Genome ID 458080) to be identical to reference sequence as reported [7]. That means, Maharashtra samples had more variations from reference sequence to begin with. But if we look and Odisha and Tamil Nadu with 30 and 31 samples accounting for 109 and 40 variations respectively, it’s evident that sequences in some regions are mutating faster than others. Another contrasting example pair is Delhi (63 samples, 54 mutations) and West Bengal (40 samples, 70 mutations). The exact mutational route can be revealed only with tracking the route of samples and spreading of infection which has not been feasible for present dataset due to paucity of information. However, we can surely say that some sequences are mutating more than the others but whether the geographical location is playing a role needs to be ascertained.

**Figure 5:**
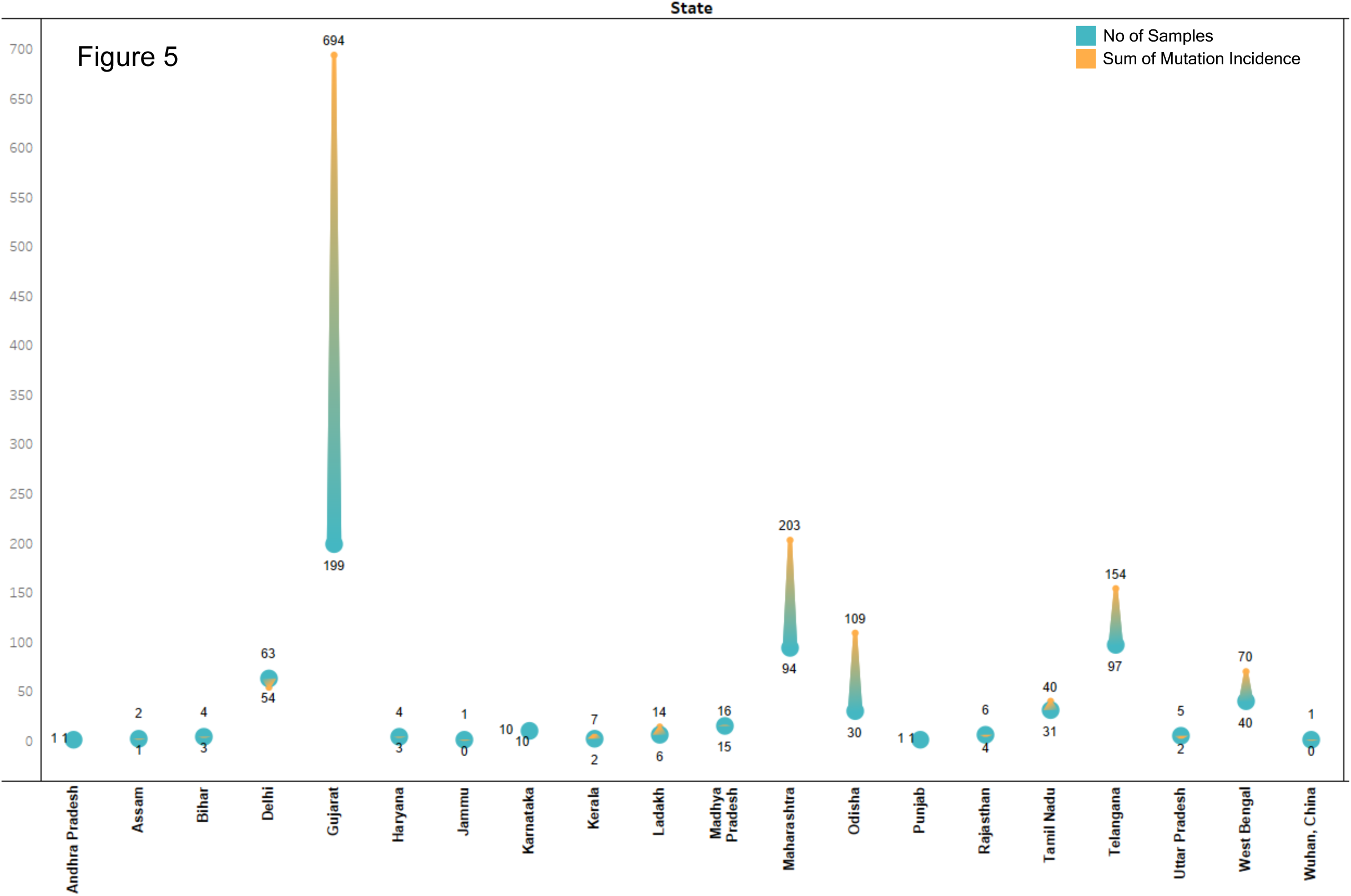
Number of samples and corresponding Sum of Mutation Incidence of SARS-CoV-2 across different states of India.

### Impact of variables sites on viral proteins

A total of 281 SNPs which were present which were altering the amino acid sequence. Their details and positions have been summarized in Table 4 and Supplementary file 5. We also ascertained the prevalence of these variants across samples. The most incident variant Q57H localized to ORF3a was present in 127 samples followed by A994D in NSP3 present in 29 samples. Amongst the silent SNPs, Y71Y in M protein was present in 117 samples followed by D294D in S protein with 69 incidences. The overall data for variants present in 10 genomes or more has been summarized in Figure 6a. Conversely, we also assessed the accumulation of variations in a given genome as summarized in Figure 6b. Interestingly, one sample (Genome ID 461495) had highest incidence of 11 mutations while 114 samples harbored just a single mutation. There were 156 samples with no mutations and 340 with more than one mutation. To account for these, the Sum of Mutation Incidence has been used in this study as explained above.

**Figure 6:**
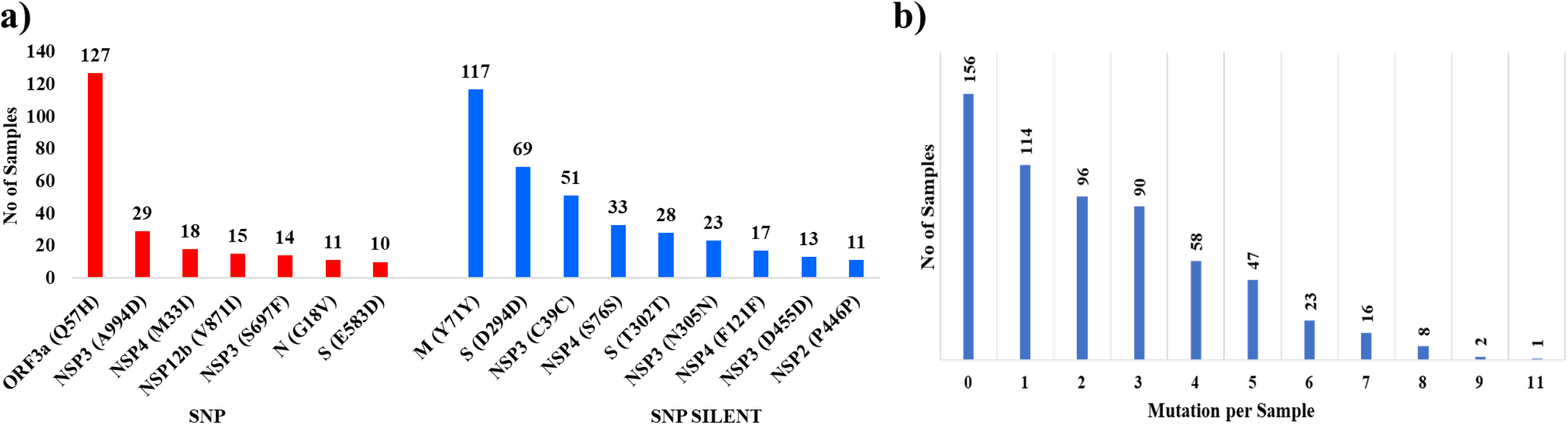
Prevalence of variant sites across studied genomes. **a) Most prevalent SNP and SNP_Silent across studied samples.** Variants which had incidence across more than ten genomes haven been represented. **b) Samples with accumulated variations in genomes**

**Table 4:** Distribution of SNPs across different proteins

| S No | Gene/ Protein | No of Variable Sites | Sum of Mutations Incidence | Mutations affecting Protein (N=N2+N3; D=D2+D3) | Neutral by Three (N3) | Disease by One/Neutral by Two (N2) | Disease by Two/Neutral by One (D2) | Disease by Three (D3) |
| --- | --- | --- | --- | --- | --- | --- | --- | --- |
| 1 | 5'UTR | 8 | 16 | - |  |  |  |  |
| 2 | ORF1a | 231 | 578 | 127 |  |  |  |  |
|  | NSP1 | 14 | 19 | 7<br>(N=6; D=1) | H81Y, G137C, D147E, V169A | R24C; V38F | R124C |  |
|  | NSP2 | 29 | 46 | 15<br>(N=14; D=1) | Y196H, L204F, K338R, I616F, Y621S, E633D, V682L, T708I, I739V, P765S | G192D; P626T; I671T; V710F | T592I |  |
|  | NSP3 | 101 | 284 | 63<br>(N=54; D=9) | A872T, T882I, Y925C, E940D, P971S, G989V, S1029I, P1054L, M1083I, H1141Y, A1268T, S1534I, T1543K, T1567I, M1588I, V1629A, S1733G, T1761I, G1861S, T1822I, T1854A, T1854I, N1871T, K1973R, S2103F, K2029E, G2035E, P2144S, L2146F, A2249V, V2372I, T2274I, T2300I, L2323V, P2480L, H2520Y, A2593V, S2625F, S2661F | D930G, D1036G, A1298V, A1649S, S1515F, A1812D, S1978P, P2046L, P2079L, S2015K, S2015R, G2118C, S2303Y, V2613F, V2629A | A1306V, P1472S, D1625V, L2146P, S2242P | G1069E, Y1675C, C2691S, A2732D |
|  | NSP4 | 26 | 109 | 14<br>(N=11; D=3) | W2769L, M2796I, H2831Y, A2994V, F3031Y, D3042N, A3143V, L3161I | T2777I, T3058I, Y3160H | L2781P, F3071Y | T3223I |
|  | NSP5 | 20 | 41 | 9<br>(N=4; D=5) | T3453A, Q3390R, P3395S | N3405L | S3386P | L3338F, A3379V, Y3472H, S3517F |
|  | NSP6 | 17 | 30 | 7<br>(N=6; D=1) | Q3826H<br>Q3826R | I3731T, P3613L,<br>D3681N, I3835T, | I3731F |  |
|  | NSP7 | 2 | 5 | 0 |  |  |  |  |
|  | NSP8 | 11 | 26 | 6<br>(N=3; D=3) | K4081R | E3962K, K4069T | M4032S, L4033F | R3993C |
|  | NSP9 | 6 | 11 | 3<br>(N=2; D=1) |  | V4181I, V4242I |  | T4249I |
|  | NSP10 | 4 | 4 | 2<br>(N=0; D=2) |  |  |  | A4271V<br>A4273V |
|  | NSP12a | 1 | 3 | 1<br>(N=1; D=0) | S4398L |  |  |  |
| **3** | **ORF1ab** | **103** | **182** | **60** |  |  |  |  |
|  | NSP12b | 43 | 84 | 22<br>(N=12; D=10) | A4487V, I4593L, S4621G,<br>A4645G, T4685I, A4798V<br>S5305L | K4451N, R4565H<br>M4588I, E4670D,<br>L4721I | K4483N, D4532G,<br>A4577V, D4676Y,<br>S4710T, M5148I,<br>V5272I | D5076G<br>G5200E<br>T4801I |
|  | NSP13 | 30 | 49 | 21<br>(N=8; D=14) | P5377S, S5490A, K5669R,<br>M5798V, V5894L I5899V,<br>T5923I | A5770V | K5364R, E5492Q,<br>I5617V, A5620S,<br>P5624L, V5820L,<br>N5827K, A5833G.<br>M5900I | W5830C<br>Y5865C<br>P5726S |
|  | NSP14 | 15 | 20 | 7<br>(N=7; D=0) | A5926S, P5971L, M5974I,<br>K6274N | L5952I, T5930A<br>S6180I |  |  |
|  | NSP15 | 8 | 17 | 5<br>(N=3; D=2) | V6518L, A6533T | G6581D |  | D6491Y<br>K6486T |
|  | NSP16 | 7 | 12 | 5<br>(N=4; D=1) | P6805S, L6909F, A6914S | K6958R |  | G6837C |
| **4** | **S** | **74** | **227** | **49**<br>**(N=36; D=13)** | L54F, N148Y, E156D,<br>A243S, S255F, G261S,<br>Q271R, T299I, T323I,<br>E471Q, A520S, T572I,<br>E583D, T602I, V622I,<br>Q677H, A706S, T761S,<br>G769V, T827I, A831S, | I434K, S494P,<br>D574Y, A892V<br>H1083Q, P1263L | T723I, F797C,<br>L828P, T941K,<br>V1068F, D1153Y,<br>C1243F | G857C<br>A930T<br>A930V<br>S1021F<br>I1179N<br>C1250F |
|  |  |  |  |  | A879S, T1027I, H1101Y,<br>V1104L, G1124V, K1181R,<br>K1191N, G1251V, Q1201K |  |  |  |
| 5 | ORF3a | 31 | 178 | 22<br>(N=9; D=13) | V13L, G18V, S74A, V77F,<br>T175I | L41F, S74F,<br>S171L, T190I | I62T, L83F, T176I | I35T, L46F,<br>L53F, Q57H,<br>C81F, L85F,<br>L86W,<br>G172C,<br>G196V,<br>G251V |
| 6 | E | 5 | 8 | 1<br>(N=0; D=1) |  |  |  | V29S |
| 7 | M | 10 | 136 | 5<br>(N=5; D=0) | D3G, A68S, H125Y | A69S, V70F |  |  |
| 8 | ORF6 | 6 | 17 | 2<br>(N=1; D=1) | I60V |  | D61L |  |
| 9 | ORF7a | 2 | 2 | 2<br>(N=0; D=2) |  |  | P45L | G38V |
| 10 | ORF7b | 1 | 1 | 0 |  |  |  |  |
| 11 | ORF8 | - | - | - |  |  |  |  |
| 12 | N | 20 | 38 | 13<br>(N=12; D=1) | S37L, L139F, A152S | P6T, P13L, G18V,<br>G30V, S33I,<br>D63N, D144Y,<br>A156S, Q160P |  | R92S |
| 13 | ORF10 | - | - | - |  |  |  |  |
| 14 | 3'UTR | 2 | 3 | 0 |  |  |  |  |
|  |  |  |  | 281 |  |  |  |  |

The impact of mutations on proteins was predicted through three different tools; SIFT, PROVEAN, ws-SNPs&GO; which classified the mutations as “Neutral/Tolerated” or “Disease/Affect Protein Function/Deleterious”. For the sake of simplicity, we have referred the results from all sites as Neutral and Disease. Though the prediction outcomes of the three tools were not in sync for all sites but since the classification of outcomes were on similar lines, the results can be represented in a binary manner with four categories. First two categories represent wherein the three tools have the same prediction; either all predicting a site to be “Neutral” or “Disease”. The other two categories represent deviation between prediction outcomes. They are “Disease by One, Neutral by Two” and “Disease by Two and Neutral by One”. For comparison between variants, any mutation predicted as Disease by Two or Three tools are considered as Disease and mutations predicted as Neutral by Two or Three tools are considered as Neutral.

The distribution of Disease and Neutral variants across the different genes of SARS-CoV-2 has been shown in Table 4 and Supplementary file 5. These could be analyzed in three aspects. First, in terms of overall incidence. The maximum variants affecting protein sequence were present in NSP3 (63) followed by Spike (S) protein with 49 variants. Secondly, if we focus only on variants with predicted outcome as “Disease” then NSP13 has a maximum of 14 such variants followed by S protein and ORF3a with 13 variants each. Thirdly, we looked at those proteins which had more Disease variants as compared to Neutral. There were five such proteins namely: NSP5, NSP10, NSP13, ORF3a, E, ORF7a. Of these NSP10 had just two variants and both of them were predicted as Disease by all three tools. Others had differential bias towards Disease variants. Thus, we can say that though some regions of the genome have more variations but mostly Neutral while others with fewer variations are more impactful in terms of their predicted impact due to more Disease variants. Conversely, mutations in some proteins can be relatively better tolerated by the viral genome.

The overall protein prediction outcomes of the 611 genomes have been summarized in Figure 7. There were total of 198 mutations (70%) and 83 mutations (30%) which are predicted to be Neutral and Disease respectively by at least two tools. These predictions suggest that even though mutations are accumulating in SARS-CoV-2, they are predominantly neutral. This is the possible reason that no major virulence or physiological deviations have been observed so far.

**Figure 7:**
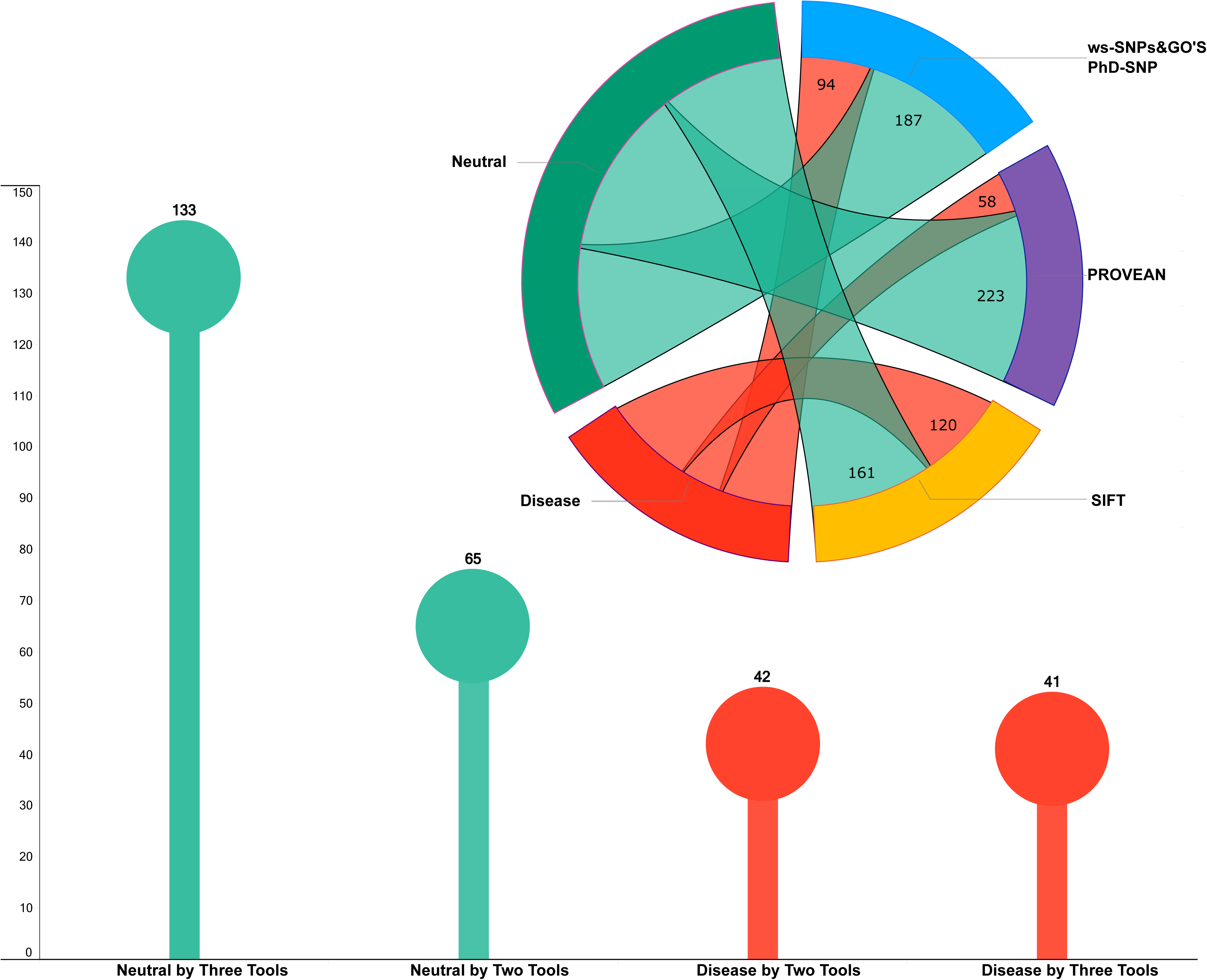
Predicting the impact of mutations affecting amino acid sequence on proteins through multiple tools.

### Mutational profile of Asymptomatic and deceased samples

In order to further assess impact of these variations we compared their prevalence across samples which were asymptomatic with those wherein the patient died. The idea was that if predictions are true, then asymptomatic samples should have more of neutral mutations whereas deceased ones should have more of disease mutations. The present congregation of samples in the study had just 2 asymptomatic samples and 15 deceased. Thereon, we included 30 new asymptomatic samples (Supplementary file 1) and compared their amino acid mutations with those of 15 deceased samples. Their comparative data has been shown in Table 5. The p value therein represents the probability that a given variant chosen at random to be Neutral or Disease. Taking the threshold as common prediction by at least two tools the data gives interesting insights. As shown and previously mentioned, for the original congregation of 611 samples, 70% mutations were Neutral (p value 0.7) and 30% were Disease (p value 0.3). The asymptomatic samples with a total of 55 SNPs affecting amino acid sequence had p value of 0.89 for Neutral variants and 0.11 for Disease variants. Corresponding data for the deceased samples with a total of 13 mutations affecting amino acid sequence had p values of 0.61 and 0.39 for Neutral and Disease variants respectively. Evidently, asymptomatic samples had majorly neutral mutations (89%) but deceased samples have a reduced share of neutral mutations (61%) and enhanced share of Disease mutations (39%). The data were analyzed in terms of p value owing to the difference in number of samples in each category. However, we do agree that a larger data set analysis for all categories with clinical correlation will provide greater insight into the impact of protein variations on SARS-CoV-2.

**Table 5:** Predicting the impact of mutations on proteins from different set of genomes

|  |  | Original congregation |  |  | Asymptomatic |  |  | Deceased |  |  |
| --- | --- | --- | --- | --- | --- | --- | --- | --- | --- | --- |
| No of Samples |  | 611 |  |  | 30 |  |  | 15 |  |  |
| Samples with no mutation |  | 156 |  |  | 0 |  |  | 01 |  |  |
| Total Variants |  | 281 |  |  | 55 |  |  | 13 |  |  |
| Mutation Type | Protein Prediction by tools | No of variants | p value | Sum of Mutation Incidence | No of variants | p value | Sum of Mutation Incidence | No of variants | p value | Sum of Mutation Incidence |
| NEUTRAL | Three Tools (N3) | 133 | 0.47 | 254 | 31 | 0.56 | 76 | 2 | 0.15 | 4 |
|  | Two Tools (N2) | 65 | 0.23 | 160 | 18 | 0.33 | 60 | 6 | 0.46 | 10 |
|  | Total = (N3+N2) | 198 | 0.7 | 414 | 49 | 0.89 | 136 | 8 | 0.61 | 14 |
| DISEASE | Three Tools (D3) | 41 | 0.15 | 187 | 2 | 0.04 | 4 | 1 | 0.08 | 2 |
|  | Two Tools (D2) | 42 | 0.15 | 70 | 4 | 0.07 | 2 | 4 | 0.31 | 6 |
|  | Total = (D3+D2) | 83 | 0.3 | 257 | 6 | 0.11 | 6 | 5 | 0.39 | 8 |
| Grand Total |  | 281 | 1 | 671 | 55 | 1 | 142 | 13 | 1 | 22 |

## Conclusions

The mutational accumulation in SARS-CoV-2 genomes is a multifactorial event with some areas of genome more prone to mutations, selective mutations being more prevalent, non-linear assimilation of mutations across various states and differential correlation between mutational impact on proteins and physiological state. Age and gender specific bias in incidence of mutations was observed. The asymptomatic samples had higher occurrence of Neutral variants while deceased samples had relatively higher incidence of Disease variants. A cross-linking of mutational dynamics and patient history will provide for better correlation and understanding of the variations in SARS-CoV-2 genomes.

## Acknowledgements

The authors thank the Department of Biological Sciences, Aliah University, Kolkata, India for all the financial and infrastructural support provided. Authors acknowledge all the authors associated with originating and submitting laboratories of the sequences from GISAID’s EpiFlu™ (www.gisaid.org) Database on which this research is based.

## Declarations

### Competing Interests

The authors declare they have no competing interests.

### Ethics approval

Not Applicable.

### Availability of Data and Materials

All data pertaining to the study has been provided as Supplementary material of the manuscript.

### Authors’ contributions

RL: Methodology, Investigation, Formal Analysis and Validation SA: Conceptualization, Supervision and Writing.

## Details of Supplementary Files

Supplementary File 1: Details of asymptomatic samples used in the study

Supplementary file 2: Localization of all the mutations observed in studied SARS-CoV-2 genomes

Supplementary files 3: Localization of variants affecting amino acid sequence in studied SARS-CoV-2 genomes

Supplementary file 4: Age, Gender and Location data for corresponding 611 SARS-CoV-2 genomes

Supplementary file 5: Details of protein prediction outcomes for 281 variants affecting amino acid sequence through different tools

## Notes

### Competing Interest Statement

The authors have declared no competing interest.

